# A hippocampal model for behavioral time acquisition and fast bidirectional replay of spatio-temporal memory sequences

**DOI:** 10.1101/343988

**Authors:** Marcelo Matheus Gauy, Johannes Lengler, Hafsteinn Einarsson, Florian Meier, Felix Weissenberger, Mehmet Fatih Yanik, Angelika Steger

## Abstract

The hippocampus is known to play a crucial role in the formation of long-term memory. For this, fast replays of previously experienced activities during sleep or after reward experiences are believed to be crucial. But how such replays are generated is still completely unclear. In this paper we propose a possible mechanism for this: we present a model that can store experienced trajectories on a behavioral timescale after a single run, and can subsequently bidirectionally replay such trajectories, thereby omitting any specifics of the previous behavior like speed, etc, but allowing repetitions of events, even with different subsequent events. Our solution builds on well-known concepts, one-shot learning and synfire chains, enhancing them by additional mechanisms using global inhibition and disinhibition. For replays our approach relies on dendritic spikes and cholinergic modulation, as supported by experimental data. We also hypothesize a functional role of disinhibition as a pacemaker during behavioral time.

## 1 Introduction

An animal’s ability to acquire and retrieve episodic memories is essential to guide its decision making and how it interacts with the environment. The hippocampus plays an important role in episodic memory encoding [1, 2, 3, 4], and in episodic memory relay into neocortex via long-term memory consolidation processes [5, 6, 7]. During active behavior (encoding phase), hippocampal neurons display asynchronous firing [8, 9], while during quiescence (sleep or quiet wakefulness), the hippocampus exhibits synchronous bursts of activity (sharp-wave ripples) [10, 11, 12], and previously stored memories are replayed on a faster timescale [10, 13, 14]. It is believed that these replays are essential in reorganizing and strengthening memory traces [15, 16]. Replays in sharp-wave ripples have also been observed to occur during wakefulness after reward experiences [13, 17, 18, 19]; in this case, replays are mostly played in the *reverse* chronological order [20, 21, 22, 19].

While it seems clear [16] that forward and reverse replays of memories during sharp-wave ripples play an essential role in memory consolidation, it is much less clear how such replays can actually be formed. Memory sequences that happen on the timescale of behavior (order of minutes) have to be stored in an online and one-shot fashion in such a way that subsequent replay is possible on a much faster timescale (order of a couple of hundred milliseconds). In addition, it is not enough to link each item in the memory sequence with its successor, as some items may occur several times in the sequence, but with different successors. As the nature of replays is best understood in the context of navigation, we next illustrate these challenges in more detail in this context.

It is known that in the hippocampus there are *place cells* which fire whenever the animal is at a specific spatial location in a given environment [23, 24, 25]. An animal which navigates in an environment, will do so varying specifics of its motion, such as its speed, the stopping locations and the duration of these stops. Spontaneous replays of such events in sharp-wave ripples, however, are independent of such speeds, stopping times or locations of the animal [13, 26, 27, 28, 29], as well during sleep [13, 17] as in quiet wakefulness [20, 30, 18, 29, 19, 31]. In addition, if the animal goes through the same location twice, turning left the first time and right the second time, the hippocampus is still able to activate the correct order of the place cells during replay (e.g., left first then right) [32].

### Our contribution

In this work, we propose a possible mechanism that addresses and resolves all challenges mentioned above. We present a model that can store experienced trajectories on a behavioral timescale after a single run, and can subsequently replay (forward and backwards) such trajectories, thereby omitting any specifics as speed, stopping times, etc. Our solution builds on two well-known concepts: (i) one-shot learning [33] and (ii) synfire chains [34]. More specifically, we make use of the fact that random connectivity is enough to learn associations between ensembles of cells in a one-shot fashion using standard forms of Hebbian plasticity. Synfire chains, on the other hand, are a well known mechanism that activates neurons in a pre-specified order. Our main contribution is an enhancement (inspired by biological observations) of the standard synfire chain approach that allows the network to alternate between an asynchronous mode, used during encoding of memories, and a synchronous mode, used during replays. We term this enhancement a **sequence encoding module**. Cells in the sequence encoding modules are called **sequence cells**. Unlike place cells, which fire at a particular location, *sequence cells* fire in a predefined position of the sequence, independent of the environment. A sequence *ensemble* consists of all sequence cells that fire at, say, the fourth position of the sequence; i.e., that correspond to the fourth layer of a synfire chain. We explain in more detail how the sequence encoding modules function in what follows. See also the video (link) which we provide in the supplementary material.

For clarity we present our approach in the context of place cell replays, even though it can be used to store any form of spatio-temporal memory sequence. Note that we require no assumptions on the geometry of the environment and on the specific trajectory taken (such as whether it contains loops). The sequence encoding modules operate in two distinct modes: an **encoding mode**, corresponding to the asynchronous firing hippocampal (theta) state observed during navigation and a **replay mode**, corresponding to sharp-wave ripples observed during sleep and quiet wakefulness. In the *encoding mode*, the synapses from sequence cells to the corresponding place cells are learned by standard Hebbian plasticity rules. For this to work, we do not need any specific (learned) connectivity between sequence cells and place cells: random connectivity with low synaptic strength between all sequence cells and place cells is enough. Associations can be learned in an online and one-shot manner between coactive cells by standard Hebbian plasticity. We use a global inhibitory population to ensure that the same sequence ensemble remains active, regardless of how much time the animal spends at a particular location. Once the animal moves to a new location, we postulate that a novelty signal activates a second group of inhibitory neurons that temporarily inhibits the global inhibitory population, thereby allowing the sequence module to move its active state to the next ensemble. We note that we do not require any specific connectivity for the inhibitory neurons (random connectivity to the sequence cells is enough). The use of disinhibitory input to shape activity patterns so they match behavioral navigation times is motivated by the experimental observation that glutamatergic interneurons in the medial septum effectively control speed-correlated firing of hippocampal neurons [35, 36]. The *replay mode*, unlike the encoding mode, exhibits extremely synchronous activity as observed in sharp-wave ripples [10, 12]. This synchrony of activity induces dendritic spikes, which produce a fast forward and/or reverse replay of the sequence cells. In replay mode, our sequence ensembles trigger, due to the learned connectivity between sequence ensembles and place cells in the encoding phase, a forward and/or reverse replay of the place cell ensembles in the correct order. Consistent with previous theoretical work [37] and experimental observations [38, 12], our model assumes that replays, dendritic spikes, sharp-wave ripples and cholinergic modulation are tightly interconnected. See the Section 2 for a more complete discussion on the assumptions of the model and how they relate to what has been observed in experiments.

## 2 Results

### 2.1 Encoding of place cell sequential activity

We demonstrate that the sequence encoding modules can account for the robust forward and reverse replays observed in the hippocampus. This is done through the simulation of a virtual mouse exploring a predefined environment. A spiking neural network of exponential integrate-and-fire neurons with physiological parameters, emulating the behavior of place and sequence cells, is associated to the mouse.

As the mouse walks through the environment, a sequence of place cells in the spiking neural network will fire, each place cell corresponding to a particular place in the environment^1^. In parallel, the sequence cells of the spiking neural network also fire, but they do so along a predefined sequential activity pattern, which is independent of the environment and the route taken by the mouse. Crucially, after exploration of the environment by the mouse, the sequence cell activity is capable of triggering replays of place cell activity, due to the plasticity of synapses between sequence and place cells. These sequence cells can be used to replay many different sequences of place cell activity. We defer the detailed explanation of the mechanism that allows sequence cells to produce the desired behavior to the next subsection. For ease of understanding we first provide some general intuition relating sequence cell dynamics to the place cell activity.

Consider as an example, a virtual mouse walking on a track, as depicted in Figure 1A. Each place cell corresponds to a specific location on the track. The place cells fire with ≈50 Hz when the mouse is at the corresponding location, and are otherwise silent. As the mouse moves along the track, a sequence of place cell ensembles is activated. The place cells are also targeted by sequence cells and plastic synaptic connections will encode the sequence of place cell activity produced by the motion of the mouse, as shown in Figure 1B. The sequence cells fire following a predefined activation pattern that possesses two modes of operation: the **encoding mode** and the **replay mode**. These correspond to the experimentally observed different modes of operation at the hippocampus during exploration and slow-wave sleep (or quiet wakefulness), respectively. In the *encoding mode*, the next ensemble in the sequence is only activated when the mouse moves to another location. The synapses from sequence cells to the corresponding place cells are learned and, thereafter, sequence cells can activate the corresponding place cell ensembles without requiring external input from the environment, as illustrated in Figure 1C. The applied learning rule is a simple Hebbian plasticity mechanism (a detailed description of the applied learning rule is given in the Section 3.2.5).

**Figure 1:**
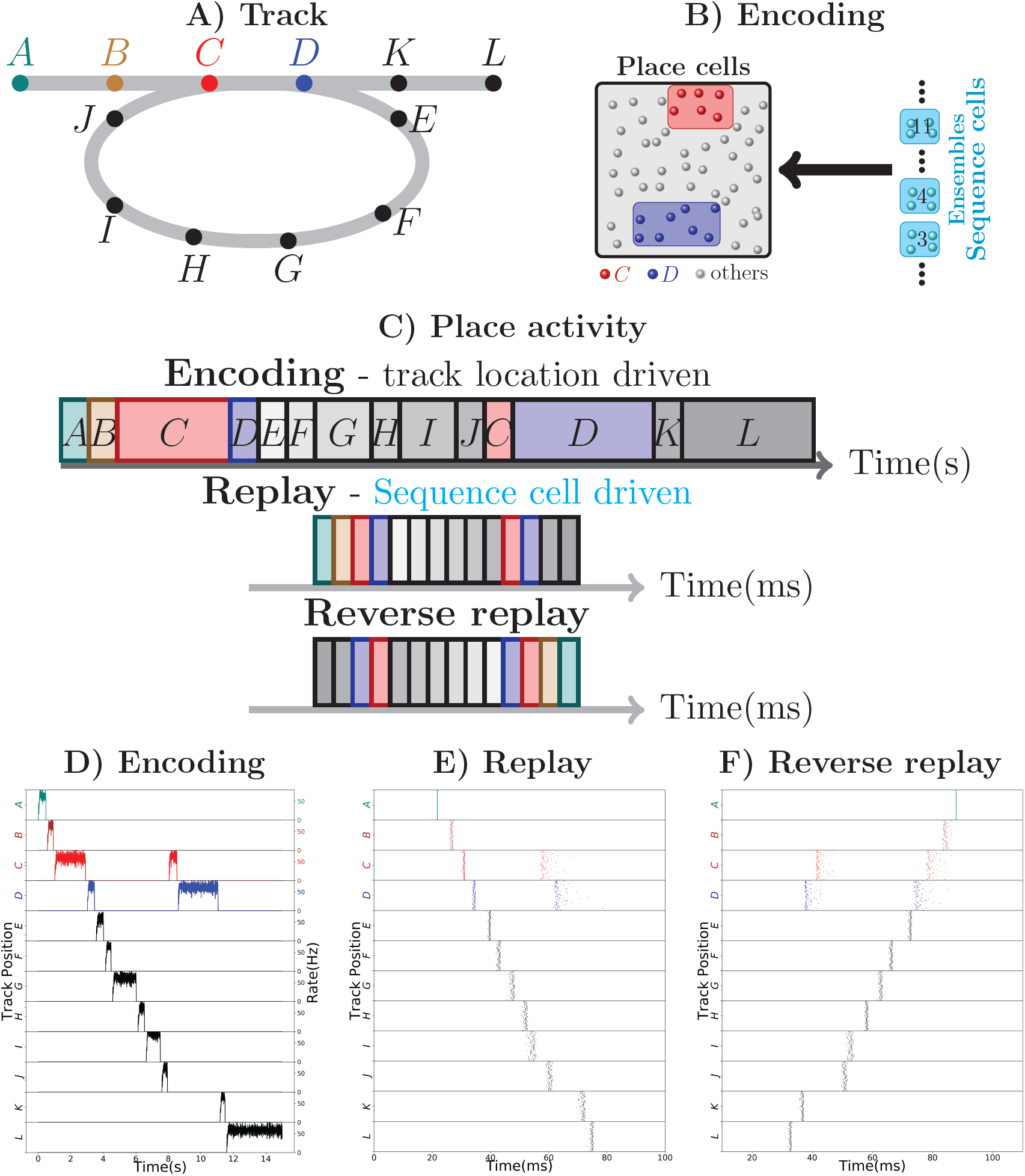
Robust replay of place cell activity. (A) An animal walks along a path, containing a loop in a 2-dimensional environment. (B) Locations in this path are associated to specific populations of CA3 place cells which fire at high rates if the animal is at said location. In this particular scenario, each position is associated with an ensemble of 100 place cells. Concomitantly, the sequence cells fire following a predefined sequential activity pattern that is explained in detail in Section 1.2. Hebbian-like synaptic plasticity connects the sequence cells to the corresponding place cell ensembles. (C) The sequence cell dynamics allow for place cell activity replay on a much shorter timescale and independent of the animal’s stopping times and locations. (D) 15 seconds simulation of place cell activity as the animal walks through the track in (A); a rate plot of the place cell ensembles corresponding to each position the animal walks through is shown; the x axis shows time in seconds, the left y axis shows the track positions corresponding to the place cell populations and the right y axis shows the firing rate of each place cell ensemble corresponding to a particular track position. Note that in addition to the loop in the path, the animal occasionally stays arbitrarily longer at particular locations. (E) Fast replay of the path taken by the animal in less than 100ms; a raster plot of place cell activity is shown; the x axis shows time in milliseconds, the left y axis shows the track positions corresponding to the place cell populations. (F) Reverse replay of the path taken by the animal; a raster plot of place cell is shown; the axis are the same as in (E). Note that each weight in the spiking network is multiplied by a gaussian multiplicative noise.

Thus, in the encoding mode, the place cell ensembles are activated in sequence as the animal moves along the track, as shown in Figure 1D. In the *replay mode*, the sequence cell ensembles are triggered one after the other, in a small time window. Through the synaptic connections that were learned in the encoding mode, the sequence cells trigger a forward replay of the place cells. A raster plot of place cell activity during the replay mode is shown in Figure 1E. Furthermore, the sequence cell ensembles can also be activated in the opposite order, and this produces reverse replay of place cell activities. A raster plot of a reverse replay of place cell activity is shown in Figure 1F. It may be noted that both forward and reverse replay are much faster than the original encoding time, that they are independent of the animal’s speed, stopping times and locations, and that the majority of place cells fire in the right order, driven by the activity of the sequence cells. Moreover, these replays can be produced after a single traversal of the path by the mouse.

We emphasize that our approach does not require any particular assumption on the geometry of the environment, the connectivity among the place cells and particularities of the path (such as loops). Sequence cells enable our model to produce replays in such a general setting, and this is discussed in the next subsection.

### 2.2 Sequence encoding modules of sequence cells

Sequence cells are organized in a specific structure that resembles a synfire chain. This structure allows sequence to produce the replays observed in place cells in Figure 1. Sequence cells might coexist in the CA3 with the place cells (or might be localized somewhere else in the hippocampus). Figure 2A shows the general structure of sequence cell connectivity, showing an example of what we term a sequence encoding module. The sequence cells can, in principle, compose many such modules that could store many parallel sequences. Each sequence encoding module consists of *ℓ* ensembles of neurons *B*_1_,…,*B_ℓ_.* Forward connections from *B_i_* to *B_i_*_+1_ are present. Backward connections, which allow reverse replays, are also present and are generally weaker than the forward connections. These ensembles share two different sources of inhibition: the **gating** inhibition *I_G_* and the **competitive** inhibition *I_C_*. The rate dynamics of individual ensembles *B_i_* have two stable fixed points, one at low firing rate (≈0Hz) and one at high firing rate (≈50Hz). Henceforth, we term the ensembles *B_i_* **bistable units**, and refer to *B_i_* as *active* at a particular time if its instantaneous rate is at the 50Hz stable point, and *inactive* if it is at the silent stable point. Furthermore, the gating inhibition *I_G_* is also either *active* (if its inhibitory neurons fire at a high rate) or *inactive* (if its inhibitory neurons are silent).

**Figure 2:**
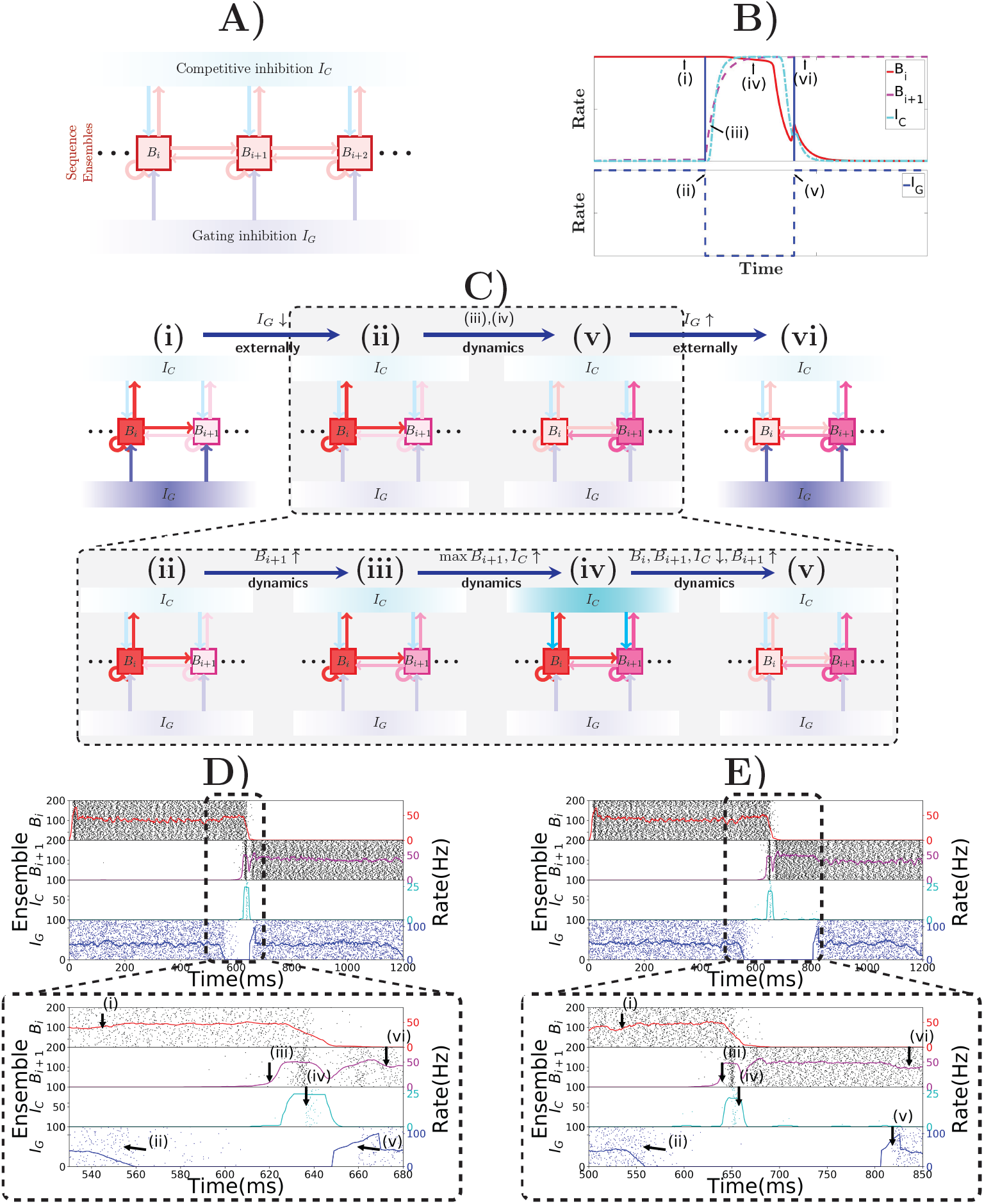
Sequence encoding modules. (A) Illustration of a sequence encoding module architecture consisting of 3 bistable units *B_i_*, *B_i_*_+1_, *B_i_*_+2_, the competitive inhibitory population *I_C_* and the gating inhibitory input *I_G_*. The inhibitory unit *I_C_* prevents two bistable units *B_j_* from having high rate at the same time, whereas the gating inhibition *I_C_* is the “break” that prevents activity from propagating from *B_j_* to *B_j_+*1. This break is externally controlled, by some mechanism that recognizes when the animal changes its location. (B) Evolution of the rates of 2 abstract rate units *B_i_*, *I_C_* and *I_G_*, disposed according to the sequence encoding architecture in (A). (C) General illustration of the states the network goes through over time, (i) and (vi) correspond to stable states in the network while *I_G_* is activated. Inactivating *I_G_* induces a dynamical process that brings the network from state (ii) into state (v). This dynamical process is depicted in further detail below with (iii) and (iv). In (iii), due to inactivation of *I_G_*, the activity in *B_i_*_+1_ picks up, driven from the activity *B_i_.* As shown in (iv), once both *B_i_* and *B_i_*_+1_ are active, they together activate *I_C_*, which gives strong inhibitory input to *B_i_* and *B_i_*_+1_ and (in (v)) forces *B_i_* to become inactive, while the recurrent activity permits *B_i_*_+_1 to recover and stay active. (D) Spiking network of sequence cells. Above, the *x*-axis shows 1.2 seconds of simulated activity. The left *y*-axis shows the indices of the cells in each sequence ensemble, and a raster plot is given. The right *y*-axis shows the rate of each sequence ensemble. Below, the period where *I_G_* is inactivated is emphasized. During this period, the stability of the system is broken and the dynamics described above take place, leading ultimately to *B_i_*_+1_ overtaking *B_i_* as the active sequence ensemble. (E) Analogous to (D), but we now purposedly enforce a longer time out of the global inhibitory mechanism. Observe that the system still functions much like in (D), so the period where *I_G_* is turned off is flexible.

As noted in the previous subsection, these sequence encoding modules possess two modes of operation: the encoding mode and the replay mode. In the encoding mode, only one bistable unit is active at a particular point in time, and this bistable unit remains active for as long as necessary, only triggering activity in the next bistable unit and turning itself inactive when the mouse moves to another location. In the replay mode, the bistable units are triggered one after the other in a very brief period of time.

The discussion of the dynamics of sequence cells is split between this subsection and the subsection 1.3. This is done to distinguish the essential building blocks of our model, presented in this subsection, from specific implementation decisions that are not essential.

#### 2.2.1 Encoding mode

The encoding mode of the sequence encoding modules relies on a specific, complex rate dynamics while the animal is exploring an environment. To describe them, and to extract the fundamental ingredients that allow a spiking neural network to produce the desired behavior, we first adopt a **mean-field approach** and collapse the bistable units *B_i_*, the gating inhibition *I_G_* and the competitive inhibition *I_C_* into abstract rate units. This approach is inspired by [39, 40]. A detailed description of this rate-based system is given in the Section 3.1.

At all times, the *competitive* inhibition *I_C_* guarantees that only one bistable unit *B_i_* can be active at any particular point in time, by realizing essentially a winner-takes-all mechanism. The *gating* inhibition *I_G_* is responsible for controlling the amount of time *B_i_* remains active. While *I_G_* is active, the currently active bistable unit remains active. If the mouse location changes, *I_G_* turns inactive, which implements a disinhibitory mechanism that allows progression of activity in the bistable units as follows (see the Section 2 for experimental evidence relating disinhibition and behavior). As soon as *I_G_* turns inactive, the active bistable unit *B_i_* drives the rate of the inactive bistable unit *B_i_*_+1_. Eventually *B_i_*_+1_ turns active and triggers the competitive inhibition *I_C_*. As *B_i_*_+1_ receives more input than any other bistable unit (if we set the connections from *B_i_* to *B_i_*_+1_ stronger than the ones from *B_i_*_+1_ to *B_i_*), it will be the only one active at the end due to the winner-take-all dynamics imposed by *I_C_*. Once *B_i_*_+1_ has taken over, the disinhibitory signal is removed and *I_G_* turns active again and prevents *B_i_*_+1_ from activating *B_i_*_+2_. Figure 2B shows this evolution of activity in a rate-based system that reproduces the desired rate dynamics.

Figure 2C illustrates the most notable states of activity the network goes through. In (i) while *I_G_* is active, *B_i_* remains active; (ii) once *I_G_* is inactive; (iii) the rate of *B_i_*_+1_ is driven up by B_i_; (iv) this, in turn, triggers an increase in the rate of *I_C_*, which (v) forces a competition among *B_i_* and *B_i_*_+1_; eventually, *B_i_*_+1_ emerges as the winner and (vi) *I_G_* is again active, preventing *B_i_*_+1_ from activating *B_i_*_+2_, while *B_i_*_+1_ stays active.

Figures 2D, E show the spiking network of sequence cells transitioning between two consecutive patterns and exhibiting the aforementioned rate dynamics. Note that the length of the period in which the gating inhibition *I_G_* is inactive is different in both. Thus, the length of the period in which the disinhibitory signal is given to *I_G_* does not need to be strictly timed. The most important requirement is that it gives enough time for the dynamics happening between (ii) and (v) to reach its stable point.

Furthermore, as will be seen in the next subsection, when the spiking network obeying these rate dynamics is presented in detail, the encoding mode has two defining characteristics other than the combined gating and competitive inhibitory processing mechanisms. They are: 1) neurons in bistable units fire *asynchronously*; and 2) transition time between one bistable unit and the next is in the order of 50 ms to 100 ms. The first property emerges from a balance of excitation and inhibition in a spiking network of neurons [41, 42] and is crucial in guaranteeing the stable rate dynamics we expect from the bistable units. The latter property is a consequence of having a spiking neural network produce the complex rate dynamics of the sequence encoding modules. Roughly speaking, for the gating inhibition *I_G_* to prevent activation of the next bistable unit, the weights between bistable units must be generally weaker than those inside each bistable unit. Since too high stable rates for a bistable unit *B_i_* are not reasonable, these weights between bistable units units will be weak. Finally, weak weights imply that some time is necessary to increase the membrane potential of the neurons beyond the spiking threshold.

In essence, in the encoding mode, the sequence encoding module activates bistable units in a sequence, with the crucial property that the sequence can be stopped, for an arbitrarily long period, at any particular point in time. This allows it to capture the essential elements of the path explored by the animal, as the sequence transitions to the next element only when a new location is reached. It may be noted that such an ability to produce compressed representations is potentially useful in other contexts. We discuss further applicability in the Section 2 of the paper.

#### 2.2.2 Replay mode

The main distinguishing feature of the encoding and replay modes is the synchrony of activity. In the replay mode, activity progresses through the bistable units in synchronous waves, reminiscent of a synfire chain [34], in a manner depicted in Figure 3A. This synchrony of activity in a bistable unit yields fast dendritic sodium spikes in the subsequent bistable unit. Dendritic sodium spikes are observed whenever, in a period of a couple milliseconds, the membrane potential at a particular branch of the dendrite of a neuron receives excitatory input above some threshold [43, 44, 45, 46]. Therefore, we assume that the neurons in bistable unit *B_i_* have the axons projecting on close branches of the dendrites of neurons in bistable unit *B_i_*_+1_. We assume a similar structure for the backward connections, as reverse replays will also be driven by dendritic spikes^2^. Whenever *B_i_* is synchronously activated, some branch of the dendritic tree of a neuron in *B_i_*_+1_ receives strong excitatory input in a few milliseconds, producing a dendritic spike that has a large depolarizing effect on the soma. Figure 3B shows the effect on the soma of one such dendritic spike, in comparison to the effect that would be produced if dendritic nonlinearities were not present.

**Figure 3:**
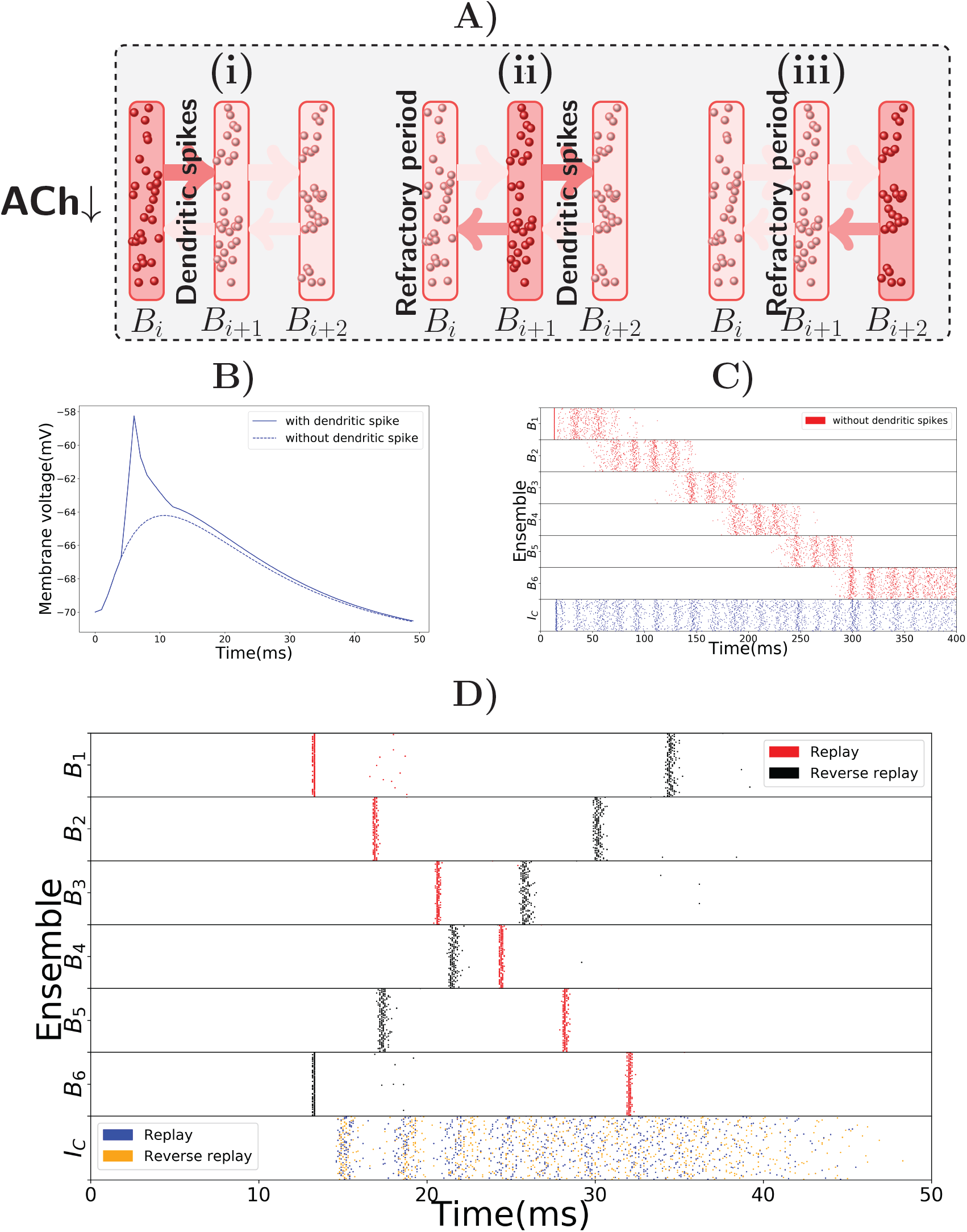
Replay mode of the sequence encoding module. (A) Synfire chain like synchronous activation of three consecutive bistable units *B*_1_, *B*_2_, *B*_3_. Synchrony triggers dendritic spikes in the next bistable unit, which in turn synchronously activate it. Activity moves in a single direction due to neurons entering a refractory period. (B) Nonlinear effect of a dendritic spike on the membrane potential at the soma of a neuron. There is a latency between the moment the dendrite triggers the dendritic spike and its effect on the soma. All dendritic spikes produce the same stereotypical current pulse effect in the soma. (C) Raster plot of the sequence encoding module in the replay mode, without any dendritic spikes. If dendritic spikes are not allowed, the sequence encoding module produces a replay of the path, but in a much slower time scale. Each transition, in the same way as during encoding, takes between 50 to 100ms. Reverse replays rely crucially on dendritic spikes and cholinergic modulatory effect and do not happen without either. (D) Raster plot of the sequence encoding module in the replay mode, now with dendritic spikes. Dendritic spikes increase the speed of replays by two orders of magnitude, making it in line with what is observed during sharp-wave ripples.

There is experimental support for the connection between synchronous events and replays, as replays are observed typically during sharp-wave ripple events, the most synchronous pattern of activity in the brain [49, 50, 12]. In our model, as shown in Figure 3C, the sequence cells can also be played asynchronously forward, without the appearance of dendritic spikes due to the lack of synchrony. In fact, this asynchronous slow replay of sequence cells produces replays that resemble what is experimentally observed during REM sleep [51]. In the model, the forward replays without synchrony are not significantly faster than the original encoding time, with transition between consecutive bistable units is between 50 ms to 100 ms. Such slow transition times are not in line with the speed of forward replays observed during sharp-wave ripples, but they are comparable to the speed of replays observed during REM sleep where sharp-wave ripples are rare and theta activity is predominant [9]. Furthermore, without using synchrony and cholinergic modulation, it is not possible to significantly reduce such transition times. If asynchronous firing of a bistable unit *B_i_*, in a couple of milliseconds, sends too large input to *B_i_*_+1_, then a simple form of inhibitory gating of activity may not prevent *B_i_*_+1_ from turning active. Even more, the non-linear effect of dendritic spikes is also crucial if the neurons in *B_i_* are synchronously activated. This is because, without the non-linear effect of dendritic spikes, it is difficult to distinguish high asynchronous firing rates, from synchronous events. Only a couple milliseconds are required for a bistable unit firing at 50 Hz to send forward the same input that it would send if it fired synchronously^3^.

Figure 3D shows a fast replay of sequence cells produced by the spiking network. Note that the transition time drops to around 5 ms, that the activity in each bistable unit is very synchronous and that most neurons spike once and in the right order.

The second distinguishing feature between the encoding and replay modes is a different cholinergic environment. In the replay mode, we assume lower levels of Acetylcholine (ACh) in the network. This is motivated by the experimental observation that replays typically occur in sharp-wave ripples during slow-wave sleep and quiet wakefulness, where lower ACh levels are documented [38, 52]. It is known that a lower ACh level makes synapses overall stronger and the network more excitable [53]. In our model, higher excitability is not necessary for forward replays, but it is strictly necessary for reverse replays. Furthermore, higher excitability facilitates faster transition times between bistable units, as well as the occurrence of dendritic spikes. In principle, if there would not be any dendritic spikes, this higher excitability of the network could decrease the transition times between bistable units, even when the bistable units fire in an asynchronous manner. Hence, the higher network excitability would allow for the bistable units to be replayed asynchronously faster than shown in Figure 3C, which would in turn allow for replays even without a synchronous mode containing dendritic spikes. However, in order for the replays to be as fast as what is observed in the literature [17, 50], the magnitude of the cholinergic effect on the network would have to be much higher than what is reported in [53]. Dendrtic spikes are therefore necessary for fast replays of activity, in line with what is reported in the literature.

In short, due to slow transition times, the encoding mode of the sequence encoding module cannot account for the fast robust replays observed in the hippocampus. The module needs another mode of activity that has faster transition times. We remark that this different mode of activity is consistent with biology, as these replays are observed during quiet wakefulness and sleep, where the hippocampus is known to be operating in a different regime. Two essential observed properties of the hippocampus in quiet wakefulness and sleep motivate the replay mode: sharp wave ripples and a different neuromodulatory environment, especially, lower levels of Acetylcholine (ACh). Evidence concerning sharp wave ripples indicate that the sequence encoding module in the replay mode might have its bistable units activated in an almost synchronous manner [12]. Lower ACh levels, as is observed during sleep and quiet wakefulness [38, 52], induce higher excitatory synaptic weights which facilitate faster transitions between bistable units.

#### 2.2.3 Reverse replays

As mentioned before, the reverse replays rely crucially on the higher network excitability produced by lower levels of ACh, whereas forward replays do not. This is because the rate dynamics of the encoding mode lead to a weight asymmetry condition. The backward connections among bistable units that enable reverse replays must necessarily be weaker than the forward connections among bistable units. In particular, this asymmetry gives a potential function for the (experimentally observed) increased excitatory weights induced by lower ACh levels [38, 52]. For the backward connections to produce synchronous activation of the bistable units in reverse order, they need to be strong enough to produce dendritic spikes. However, as they are weaker than the forward connections, in the encoding mode, asynchronous activation of a bistable unit should not produce dendritic spikes in the next one. If the excitatory synaptic weights are not generally increased in the replay mode it is difficult to have, at the same time, dendritic spikes at the backward connections in the replay mode and a fully functioning encoding mode. One either gets instability in the encoding mode, or not enough dendritic spikes in the replay mode. Therefore, reverse replays are made possible by the difference in the neuromodulatory environment, in addition to sharp-wave like synchronous activity triggering dendritic spikes.

Observe, additionally, that dendritic spikes have a refractory period of about 5 ms [47, 48, 37] and that within this refractory period the respective dendrites do not transmit any spikes. It is this refractory period that forces the synchronous wave to travel in a single direction. This is also shown in Figure 3A. In this case, if bistable unit *B_i_* is synchronously activated by dendritic spikes from *B_i_*_−1_ (from *B_i_*_+1_), it will produce dendritic spikes only in *B_i_*_+1_ (in *B_i_*_−1_), as the neurons of *B_i_*_−1_ (of *B_i_*_+1_) are in the refractory period, respectively. Moreover, Figure 3D shows a reverse replay of activity produced by the spiking network.

In essence, the asynchronous encoding mode has a preferred direction for the activity to travel, as the forward connections between bistable units *B_i_* and *B_i_*_+1_ are stronger than the backward connections between *B_i_* and *B_i_*_−1_. Dendritic spikes, cholinergic modulation and the refractory period of neurons allow the sequence cells to be activated in both directions, depending on which bistable unit the synchronous volley starts.

### 2.3 Sequence encoding modules - spiking network

In the previous subsection, the essential building blocks of sequence encoding modules were discussed on a conceptual level. In this subsection, we describe more detailed conditions that are necessary so the dynamical system comprising our spiking network model behaves as intended.

#### 2.3.1 Bistable units

In the spiking network, each bistable unit consists of a recurrently connected population of excitatory and inhibitory cells as depicted in Figure 4A. To produce bistable units where the rate dynamics maintain two stable fixed points, one around 0 Hz and one around 50 Hz, we analyse the gain function [54] of excitatory and inhibitory neurons as shown in the left of Figure 4B. There, the rate of excitation (*red*) and inhibition (*teal*) is shown for varying recurrent excitatory (inhibitory) and compared with the identity (*black*). Formally, to obtain the stable points of activity one has to solve the system of equations: *r_E_* = *F*(*r_E_*, *r_I_*) and *r_I_* = *G*(*r_E_, r_I_*), where *F* and *G* represent the evolution of the rate of excitatory and inhibitory neurons when receiving excitatory rate *r_E_* and inhibitory rate *r_I_*. For the plot, the implicit solution *r_I_*(*x*) = *G*(*x*, *r_I_*(*x*)) is approximated for all *x*. For the excitatory gain function, the function *F*(*x*, *r_I_*(*x*)) (in *red*) is plotted and compared to the identity (in *black*). An analogous procedure is done to plot the inhibitory gain function *G*(*r_E_*(*x*), *x*), where we now define *r_E_*(*x*) implicitly via the equation *r_E_*(*x*) = *F*(*r_E_*(*x*), *x*).

**Figure 4:**
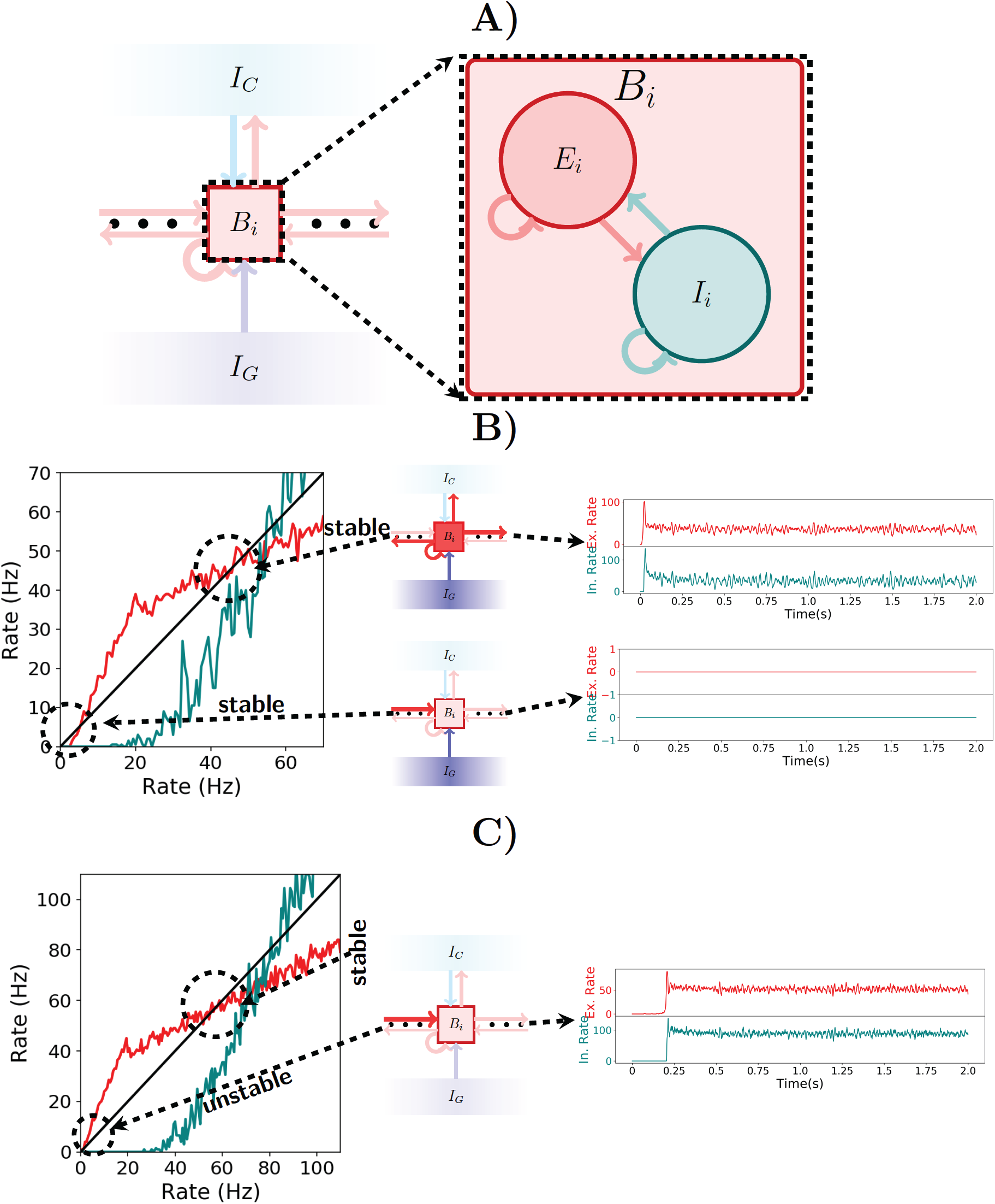
Spiking network. (A) Illustration of a bistable unit consisting of both an excitatory and an inhibitory population. (B) On the left, the gain function of excitatory (red) and inhibitory (teal) neurons inside a bistable unit *B_i_* is depicted. As shown in the center, if *I_G_* is turned on, then the shape of the gain function produces two stable states, one at 0Hz and another at approximately 50Hz. The stable states are shown on the right. (C) Analogous to (B), but now with *I_G_* turned off. As seen on the left, the gain function shape changes when receiving input from the previous pattern and the stable state at 0Hz ceases to exist. The network always settles for the only stable point around 50Hz, a fact which is shown on the right.

Note that the excitatory gain function is below the identy for low rates in Figure 4B but not in Figure 4C. This implies that there is a stable point at 0 Hz while the gating inhibition *I_G_* is active, even when receiving input from the previous bistable unit, but this stable point disappears when *I_G_* is turned inactive. This allows the activity to move forward. On the other hand, in both Figure 4B and 4C, the excitatory gain function is above and then below the identity around the corresponding stable points. This characterizes the stable point as well as the stable region around it.

#### 2.3.2 Encoding mode

The spiking network operates in the encoding mode if the initial excitatory input that activates the first bistable unit does so in an asynchronous manner. If the bistable units exhibit the correct rate dynamics, then the gating and competitive inhibitory mechanisms guarantee that it behaves as intended, again, by a through analysis of the gain functions of excitatory and inhibitory neurons. From the conditions given by the rate dynamics we have: (1) if the gating inhibition *I_G_* is inactive, then the input sent by *B_i_* to *B_i_*_+1_ should be enough to activate *B_i_*_+1_; (2) if *I_G_* is active, then *B_i_* remains active; and (3) while *I_G_* is active, *B_i_*_+1_ remains silent. Figures 4B, C explain the functioning of the gating inhibitory mechanism. In Figure 4B observe that if *I_G_* is activated then the bistable unit *B_i_* maintains both high and low rate stable points, even when receiving input from the previous bistable unit.

The behavior of the network changes once *I_G_* is inactivated. If bistable unit *B_i_* receives input from the previous bistable unit then the stable point at 0 Hz disappears and only the 50 Hz stable point remains. This is shown in Figure 4C. Observe that if the gain functions behave as shown in Figure 4B and Figure 4C then the spiking network will satisfy conditions (1), (2) and (3) from above.

We observe a few additional points. Increasing the rate of *I_G_* decreases the gain functions of excitatory and inhibitory neurons. This enlarges the stable region around 0 Hz, but shrinks the stable region around 50 Hz, and also shifting to a slightly smaller rate value. Thus, satisfying the three conditions, corresponds to not shrinking the stable region around 50 Hz by too much in (2) and enlarging the stable region around 0 Hz enough to satisfy (3). This observation also implies that it is not feasible to have the high firing rate stable point at too small firing rates, as satisfying all three conditions is difficult.

Again, from the rate dynamics we see that: once *I_G_* is inactive, and *B_i_*_+1_ gets active, the competitive inhibition *I_C_* will spike and guarantee that only *B_i_*_+1_ remains active. The competitive inhibition *I_C_* functions here as a coincidence detector of two active bistable units. It remains inactive if only a single bistable unit is active, but once two or more are active, its neurons receive enough excitatory input to spike. Such behavior is a simple consequence of the thresholding of the membrane potential of neurons to produce an action potential. With such a one spike volley, *I_C_* gives strong enough inhibitory input to quickly inactivate *B_i_*, but *B_i_*_+1_ recovers and keeps active as it gets excitatory input from *B_i_* and from itself. As soon as the activity *B_i_*_+1_ stabilizes, the gating inhibition *I_G_* turns active and prevents activity from going onwards to activate *B_i_*_+2_.

#### 2.3.3 Replay mode

The spiking network is in the replay mode if the first bistable unit is activated in a synchronous manner. This synchronous activation of the neurons in a bistable unit triggers dendritic spikes in neurons in the next bistable unit, which in turn, trigger synchronous spikes at the corresponding neurons and the activity can continue in a synchronous wave, much like in a synfire chain [34]. Such activity travels only forward because the dendrites have a refractory period of 5 ms after transmitting a dendritic spike to the soma.

#### 2.3.4 Reverse replay

The replay mode of the spiking network can produce reverse replays if the last bistable unit is activated synchronously. As in the forward replays, the activity proceeds synchronously through dendritic spikes. Again, it has a single direction of motion, due to the refractory period of the dendritic compartments that produce dendritic spikes. We emphasize that the increased synaptic weights in the replay mode, due to lower ACh levels, are essential if one wants the encoding and reverse replay modes to reliably behave as intended.

### 2.4 Robustness to noise and parameter variations

We want to point out that the chosen parameters are biologically plausible and do not need to be fine-tuned. To this end, we produce Figure 5, showing that the choice network ensemble sizes and the choice of network weights are not crucial for the spiking network to behave as intended. In Figure 5A and Figure 5B we double the network ensemble sizes, while halving all the weight parameters (alternatively, one could halve the connection probability). The encoding and replay modes function the same way as before. In Figure 5C and Figure 5D we change all the used parameters following the guide of the gain functions and show another set of parameters where the encoding and replay modes behave as intended. We observe that while the conditions of the dynamical system imply one cannot take a single parameter and change it significantly, one can change the parameters if a certain set of reasonable conditions is ensured (in other words, one has to respect the coupling among different parameters).

**Figure 5:**
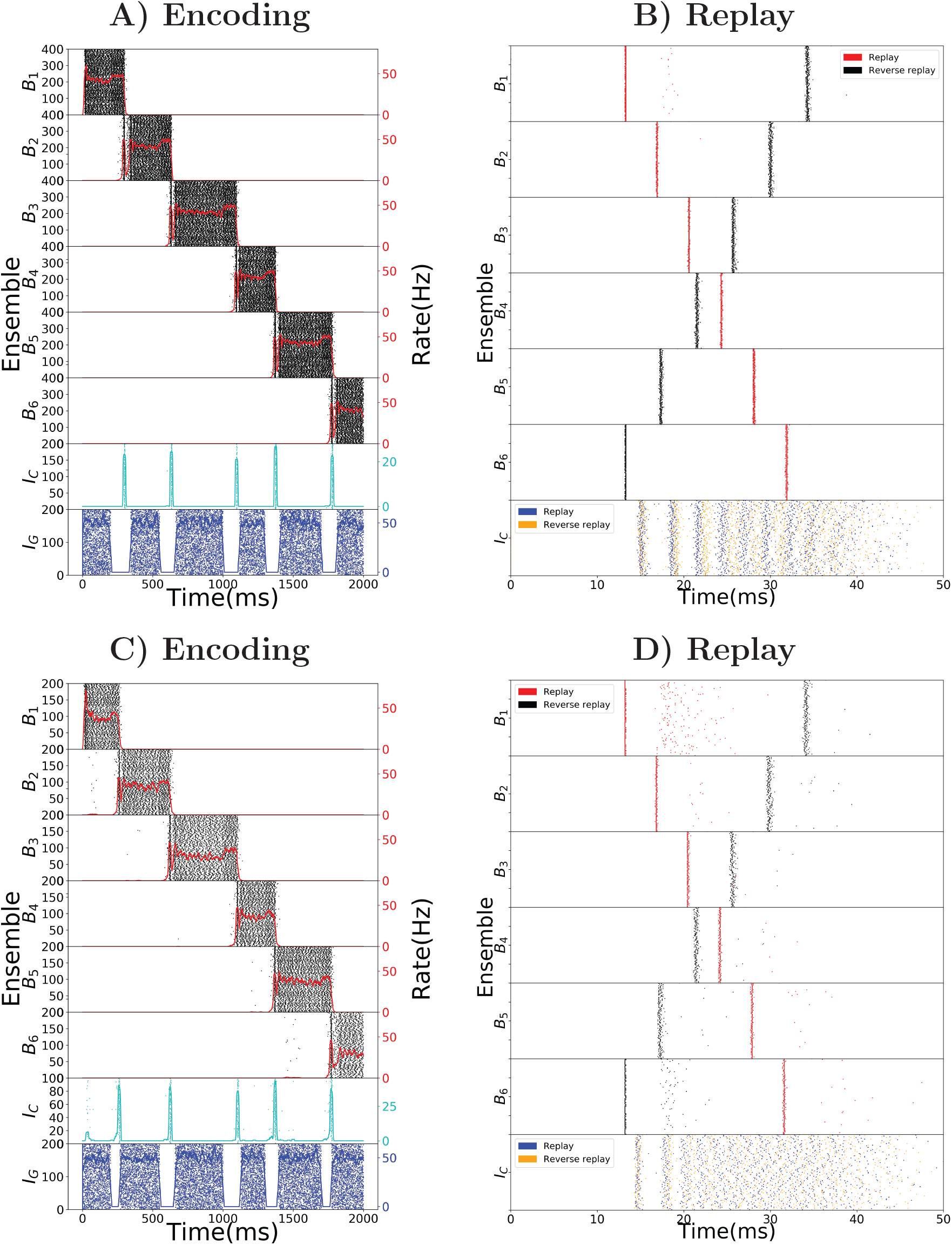
Robustness to parameter variations. (A) Raster plot of the encoding mode of the sequence encoding module with 6 patterns. The size of ensembles *B_i_* and *I_C_* are doubled, while the connection probability is halved (alternatively, one can half all corresponding weight parameters). This shows that the size of each ensemble is not important for the model to work. (B) Analogous to (A) but now the replay mode is shown. Again each ensemble size is doubled. (C) Raster plot of the encoding mode of the sequence encoding module, with an alternative parameter set (see Table 6). This shows that the choice of parameters we used is not unique. Instead, as long as the conditions on the gain functions are satisfied, there is a large range of weight parameters where the model performs as expected. (D) Raster plot of the replay mode using the alternative parameter set (see Table 6).

As a final note, we emphasize that the plots in Figure 1 were produced using a gaussian multiplicative noise on each individual weight. It is not crucial for the spiking network that each individual weight is as defined, only that the total average weight inside a particular ensemble or between ensembles is close to the desired values.

## 3 Discussion

### Summary

In this work, we show a model which can store in behavioral time place cell activity traces corresponding to paths run by an animal and produce fast bidirectional replays of these place cell activity traces during sleep and quiet wakefulness, even after a single run of the path. Furthermore, our approach works independently of the geometry of the environment and of particularities of the path taken (such as loops present in it). As observed in the literature, the replays produced by our model have constant speed and are not influenced by the potentially varying speed of the mouse during the run. As the primary conclusion of our model, we predict the existence of a population of sequence cells, dedicated to the task of encoding of sequential activity, which enables the model to properly deal with loops in the path of the mouse. This is an experimentally verifiable hypothesis, as sequence cell firing should be distinguishable from place cell firing, as well as from head direction cells, border cells and so on (time cells [55] could pose a challenge as sequence cells also measure time in a certain sense). Sequence cells enable behavioral time storage of paths traversed and subsequent spontaneous fast replay of place cell activities as observed in rodents during slow-wave sleep and quiet wakefulness. The sequence cells are organized in sequence encoding modules where, crucially, the activity progresses through stereotypical sequences of ensembles. The sequence encoding modules possess two modes of operation one used during encoding and another during replay. The encoding mode relies on a disinhibitory mechanism to control the speed of asynchronous activity progression in the ensembles. The replay mode relies on synchronous activity and resulting dendritic spikes to trigger fast replay of the sequence encoding modules. Lower levels of ACh, leading to higher excitability, allow the sequence encoding modules to be played in reverse - which would also account for the observed reverse replays of place cell activity in rodents. In addition, reverse replays happen mostly after rewards, which in our model, permit the sequence to be played in reverse to find the causes that produced the reward. In particular, a very long sequence encoding module can be used to store multiple paths, by just using the reverse replays to mark the relevant starting and ending positions in the sequence. Effectively, this allows the sequence to be split up in many subsequences, each corresponding to a relevant memory.

In the upcoming subsections, we discuss the bioplausibility of the properties we used, such as how the sequence encoding modules can be formed, the form of inhibitory processing, the dendritic spikes and the different cholinergic environment in movement and quiescence.

### Formation of sequence encoding modules

The sequence encoding modules proposed here rely on complex dynamics and a specific connectivity structure. Their structure can either be hardwired, as a product of evolution, or perhaps more likely, emerge as a consequence of the interaction of the many plasticity rules present in the hippocampus. For example, sequences of ensembles forming long synfire chains can spontaneously emerge as a product of spike-timing dependent plasticity (STDP) and axon remodeling [56], or a three factor variant of STDP [57] which is modulated by global population activity [58]. Such long sequences of ensembles form the basis of the sequence encoding modules in here. In addition, in [59], experimental evidence is presented for attractor dynamics in the hippocampus. Their data shows that linear changes in the environment context induce dynamic, non-linear changes in hippocampal activity patterns. Moreover, [60] present evidence for autoassociative and heteroassociative dynamics in the hippocampus. Densely connected populations representing attractor states (autoassociative dynamics) are sparsely connected among themselves (heteroassociative dynamics), a characteristic which permits sequential processing of information. Their data reveals a fundamental discretization in the memory retrieval processes of the hippocampus, supporting the idea that sequences of attractor states are sparsely connected, as is the case in the sequence encoding modules.

### Inhibitory neural processing

Crucial to the proper functioning of the encoding mode of the sequence encoding modules are the two forms of inhibitory neural processing, the gating inhibition and the competitive inhibition. Speed-correlated firing of hippocampal neurons has been reported and is controlled by glutamatergic interneurons in the medial septum [35, 36]. The firing rates of these glutamatergic interneurons are proportional to the speed of locomotion and control the theta oscillations during locomotion, being crucial to the initiation of theta oscillations. These glutamatergic septo-hippocampal projections perform disinhibitory control of the activity in areas CA3 and CA1, which is in line with our proposal of a gating inhibitory mechanism that is switched off during movement, possibly via some disinhibitory mechanism.

Furthermore, a specific type of inhibitory interneurons, called VIP interneurons, are known to have a disinhibitory function and seem to be ubiquitous in the brain [61, 62, 63]. These VIP interneurons perform disinhibitory control in the brain which is activated under specific behavioral conditions [63], a fact which is consistent with our hypothesis that the gating inhibition is switched off when the animal changes its location.

The winner-take-all dynamics implemented using the competitive inhibition are commonplace in the theoretical neuroscience literature [64, 65]. Furthermore, anatomical studies have shown that cortical networks contain essential features which can be implemented via winner-take-all circuits [66, 67].

As the above examples show, there exists a large diversity of inhibitory interneuron subtypes each potentially serving a different purpose [68]. Such diversity of interneuron types and functions is a strong indicator that the simple forms of inhibitory control proposed here can be realized in the brain.

### Sharp-wave ripples, dendritic spikes and replays

Previous works [37] also suggest the relation between dendritic spikes, sharp-wave ripples and replays. Our work goes along similar lines, suggesting that replays are a product of synfire chain like behavior, driven by dendritic spikes, of special populations of neurons, that fire in a stereotypical manner, independent of instantial specifics of an animal’s movement. Sharp-wave ripples are the most synchronous events reported in the hippocampus [69]. The replay of place cell activities is known to be concurrent with ripple events [26, 17]. Such synchronous events favour the occurrence of dendritic spikes, which, in turn, favour synchronous excitatory activity along a feedforward structure [37, 70].

### ACh levels and neuromodulators

It is known that Acetylcholine levels are significantly reduced in slow-wave sleep and quiet wakefulness, in comparison to an awake state [71, 72, 38]. It is further believed that such change in the cholinergic modulation is crucial for the process of acquiring and consolidating new memories [73, 38, 52]. In accordance with this work, [52] proposes that different levels of Acetylcholine during exploration, slow-wave sleep and quiet wakefulness set the appropriate dynamics of the network for encoding and consolidating memories (replays). In fact, septal cholinergic activation to CA3 pyramidal neurons has been experimentally shown to enhance theta oscillations and effectively suppress sharp-wave ripples in the CA3 network [38, 52, 74]. The nicotinic receptors either on the soma, pre-synaptic boutons, or post-synaptic spines of CA3 pyramidal cells can effectively inhibit firing of CA3 pyramidal neurons. Another potential inhibitory mechanism could be through the GABA-B receptors on CA3 pyramidal neurons, which can serve a similar role, although the origin of inputs to GABA-B receptors are currently unknown.

Lastly, a recent study [75] indicates that replays, in both directions, are more likely to occur after rewards are obtained. In particular, their results suggest that reverse replays could be a mechanism to reinforce experiences that produced rewards. This connects nicely to our work, as replays (and especially reverse replays) in our model rely on higher excitability of the network which could be further provided by increased dopamine effects.

### Other models of sequence replay

There are alternative models that achieve partially similar results to the ones reported here. These previous works fail to produce replays which cope with the irregularity of behavior and need specific assumptions about the geometry of the environment or about the trajectory taken (e.g. that it does not contain loops). In [76], a short-term synaptic depression mechanism is shown to produce two different regimes of activity of place cell rate dynamics, that can account for replays observed in the literature [13, 17] and preplays [77, 78, 79], at least under the assumption that a map of the environment is stored in the synaptic connections among the place cell rate units. In [80], it is studied how standard plasticity rules [81, 82] shape the patterns of activity of place cell rate units and it is found that the theta rhythm speeds up encoding of newly explored environments. The mechanism for generating the replays is the same as the one used in [76]. In [83], a spiking network consisting of Hebbian assemblies that are weak and sparsely connected uses pre-existing recurrent connections to quickly amplify weak signals. As in our work, the assemblies are activated like a synfire chain during replays and network states are altered by globally changing neuronal excitability (say, by cholinergic modulation). The authors of [37], in line with our work, propose that sharp-wave ripples and replays are interconnected and that nonlinear dendritic computation, in the form of dendritic sodium spikes, accounts for the observed replays in the rodent hippocampus. As in this work, the dendritic spikes distinguish between an asynchronous state present while the animal is awake and a synchronous activity state, related to sharp-wave ripples, occurring during slow-wave sleep and quiet wakefulness. In [70], dendritic spikes are again used in the generation of replays and preplays of activity. In an hypothesis similar to ours, CA3 possesses innate sequential structure that gives rise to spontaneous firing sequences and can be used to one-shot learn place fields. Still, we note that our sequence encoding modules are the first to account for a compressed representation of place cell activity to produce replays, both forward and reverse, which are independent of the instantial specifics of the animal’s movement and environment.

### Compressing sequence representations in other contexts

In this work, we restrict sequence processing to a navigation task because the hippocampus is extensively studied in this regard and there are many reported phenomena (sharp-wave ripples, cholinergic modulation, forward and reverse replays, dendritic spikes) which fit well into our framework of sequence encoding modules.

Still, the ability to divide complex tasks into simpler, compounding, sequential subtasks is essential to understand and perform the original more complex task. Identifying these subtasks and their interconnectedness, compressing useful sequences into the most relevant parts and factoring out the non-relevant ones, is an important characteristic if one wants to learn ever more complex behavioral patterns. It is a given that animals perform this form of sequence processing operation on a high-level. In this work, it is further argued that this sequence processing is such an essential operation, that the brain might develop special modules of cells capable of extracting the relevant subparts contained in a more complex task.

## 4 Materials and Methods

### 4.1 Reduced rate encoding model

To understand the dynamics of the spiking network, we consider a reduced model where we collapse ensembles into abstract rate units. As before, we denote by *B*_1_,…, *B_ℓ_* the *ℓ* bistable units that form the sequence, by *I_C_* the competitive inhibitory population to all *ℓ* ensembles and by *I_G_* the gating inhibitory input. The activity of unit *X* at time t is given by λ*_X_*(*t*) which corresponds to the rate of the ensemble. The units are connected into a network which dictates their interaction. We denote the interaction from unit *X* to itself by *w_X_*, and the interaction strength from unit *Y* to *X* by *w_XY_*. The connectivity is as in Figure 2A. We assume that interactions with an inhibitory (excitatory) source negatively (positively) affect the target’s rate. Moreover, we assume that the activity 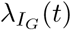 alternates between high and low rate states as depicted in Figure 2B,C. The rate of other units evolves according to the following set of differential equations:

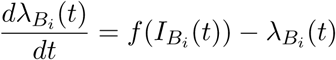

for the rate of unit *B_i_* and

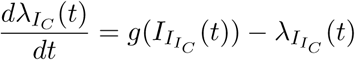

for the rate of unit *I_C_*, where 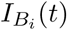 is:

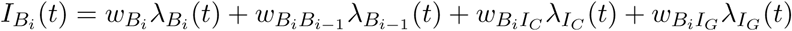

and 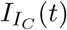 is:

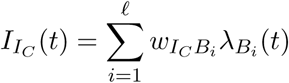

and the functions *f* and *g* are generally chosen as saturating functions such as the sigmoid function 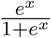. These equations represent how the rate of the populations in the spiking network should evolve, with *f* and *g* being gain functions of neurons and 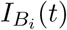 and 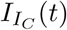 being the total input received by the respective neurons. The goal is to set the parameters of the system such that the activity evolves as in Figure 2C.

To produce the dynamics of Figures 2B,C, it is enough to study this rate model with two bistable units *B_i_* and *B_i_*_+1_, with the additional constraint that weights from and to *B_i_* are equal to those from and to *B_i_*_+1_. This allows for a simple extension to *ℓ* bistable units, as the symmetry of the weights guarantees that the system will have the same rate dynamics independent of which bistable unit is currently active. In particular, with only two bistable units, it is possible, under the assumption that *f* and *g* are piecewise linear, to write a simple dynamical system of equations that can be fully solved and understood.

Moreover, it is crucial that *f* is chosen to guarantee bistable rate dynamics of each *B_i_*, that is, specifically that close to zero or negative input produces silent (zero rate) responses and that large enough inputs produce a maximal rate response. In addition, from *g* it is required that it guarantees the behavior we expect from *I_C_*, namely, that it serves as a detector for two bistable units being active at the same time, producing maximal response if it receives input from two bistable units close to the maximal response, and being silent if it receives not much more than what a single maximally active bistable unit can produce.

Given this, one can state simple requirements that are essential: (1) that the system is at a stable point while *B*_1_ is at its maximal rate, *B*_2_ silent and *I_G_* is at the high rate state. (2) that *B*_1_ can activate *B*_2_ to its maximal rate once *I_G_* is at the low rate state; (3) that the rate of *I_C_* increases when both bistable units are jointly active; (4) *I_C_* being active is strong enough to turn silent both *B*_1_ and *B*_2_; (5) the speed of the inhibitory *I_C_* dynamics is faster than that of *B*_1_ and *B*_2_, which can force *B*_1_ below the point where it can reactivate itself, while *B*_2_ remains above said point. In particular, the speed of inhibitory *I_C_* dynamics is crucial to guarantee that the rate system can produce the desired rate dynamics with symmetric weights.

These requirements of the rate-based system, therefore, imply that one of three possible outcomes can happen, where only the third is desired: (1) *B_i_* and *B_i_*_+1_ turn both inactive; (2) the rates of *B_i_*, *B_i_*_+1_ and *I_C_* oscillate; (3) *B_i_* turns inactive, but *B_i_*_+1_ stays active, driven by activity in *B_i_* and by its own dynamics.

### Overview of parameters - rate model

Table 1, we summarize the notation and parameters used in the rate reduced model.

### 4.2 Spiking network

#### 4.2.1 Neuron models

In the spiking version of the model each ensemble unit is composed of an excitatory (*E_x_*) population with *N^E^* units and an inhibitory (*I_x_*) population with *N^I^* units where the lowercase subscript *x* corresponds to the index of the pattern. We follow the convention of denoting the populations by superscript and the neuron index by subscripts. The membrane potential dynamics of neuron *i* in population *X* follow:

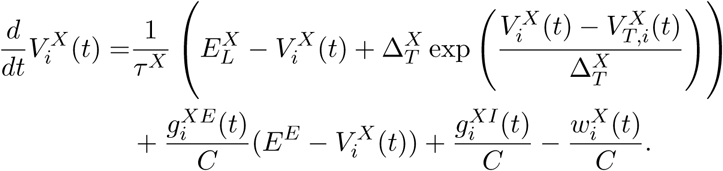

**Table 1:**
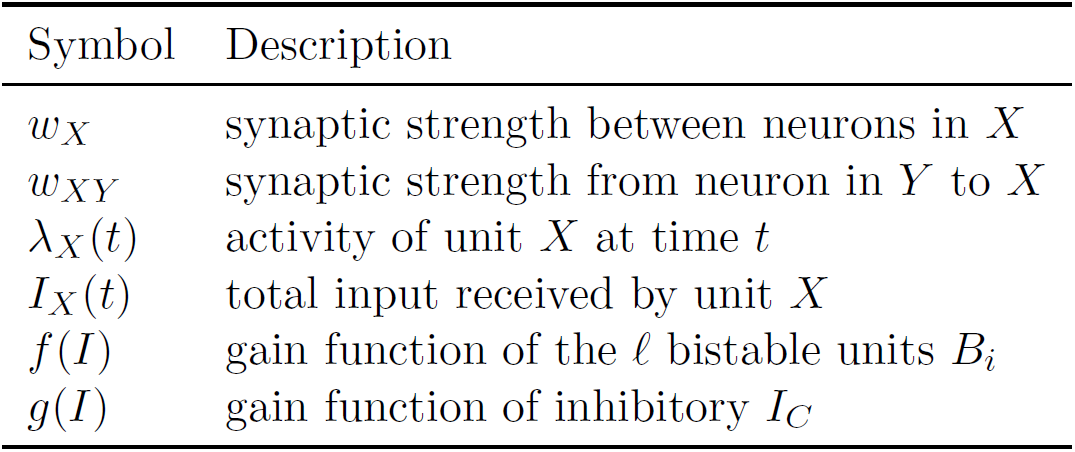
Summary of parameters for the rate model.

| Symbol | Description |
| --- | --- |
| $w_X$ | synaptic strength between neurons in $X$ |
| $w_{XY}$ | synaptic strength from neuron in $Y$ to $X$ |
| $\lambda_X(t)$ | activity of unit $X$ at time $t$ |
| $I_X(t)$ | total input received by unit $X$ |
| $f(I)$ | gain function of the $\ell$ bistable units $B_i$ |
| $g(I)$ | gain function of inhibitory $I_C$ |

The parameter values are summarized in Table 2 and the dynamics are consistent with [84, 39]. The main difference between the excitatory and inhibitory units is that the excitatory neurons have an adaptation current and an adaptive threshold [85, 86] whereas the inhibitory neurons are non-adapting. The adaptive threshold follows:

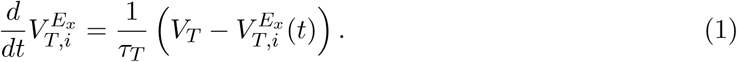

We record a spike when the voltage exceeds 20mV and we reset it to and hold it at *V*_re_ for the refractory period duration *τ*_abs_. For an adaptive threshold we set it to *V_T_* + *A_T_* after a spike. The adaptation current for a neuron in population *E_x_* follows:

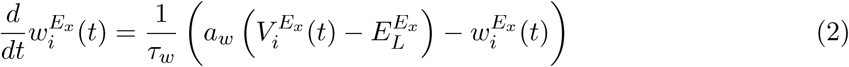

and it is increased by *b_w_* when neuron *i* spikes.

#### 4.2.2 Synapse dynamics

Connection probability between population *X* and *Y* is given by *p^XY^* and the strength of a connection between unit *i* ∈ *X* and unit *j* ∈ *Y* is given by 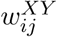. The total conductance of neuron *i* in population *X* from a source of type *Y* (excitatory or inhibitory) follows:

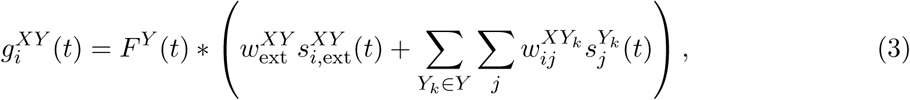

where 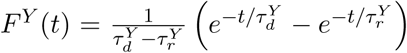 is the synaptic kernel for input from neurons of type *Y*, *Y_k_* corresponds to subpopulations of type *Y*, * is the convolution operator and 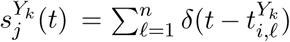 denotes the spike train of neuron j in population *Y_k_*. Additionally, each neuron receives external input given by the spike train 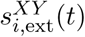 which follows a homogeneous Poisson process with rate 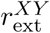 and weight 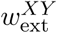 for neurons in population *X* where *Y* can either be an excitatory (*E*) or an inhibitory (*I*) source. The resolution of the simulation is 0.1ms and the coupling parameters can be found in Table 3.

#### 4.2.3 Dendritic spikes

Dendritic spikes are triggered by excitatory spikes arriving in a short time span at a particular position in the dendrite of a neuron. We incorporate them by setting the synaptic conductance of the afferent input to produce a stereotypical current pulse, using parameters observed in single-neuron experiments [43, 44, 45, 46] and similar to the one used in recent spiking network models [47, 48, 37]. Whenever the excitatory input to a (dendritic branch of) a neuron changes the conductance of the neuron by a threshold *θ_dspike_* in a time window of 2 ms, a dendritic spike is initiated. This produces a stereotypical current pulse at the soma whose effect is triggered with a time lag of τ*_DS_* after the dendritic threshold is crossed. This time lag is on the same order as the one observed in single neuron experiments. The stereotypical current pulse, like that of [37], is described by:

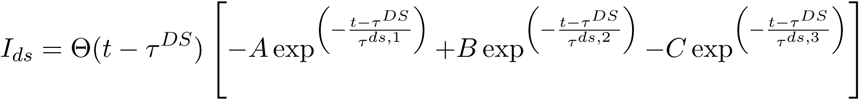

where the factors *A, B, C* and the decay time constants *τ^ds,^*^1^, *τ^ds,^*^2^, *τ^ds,^*^3^ are chosen so that the depolarizing effect on the soma fits experimental data [43, 44, 45, 46], and whose values are shown in Table 4. Moreover, we add a refractory period *t^ref,ds^* = 5 ms, as in [47, 48, 37], to these dendritic branches after the stereotypical current pulse has influenced the soma, in accordance to the saturating effect of somatic depolarization of dendritic spikes reported in the literature [43].

#### 4.2.4 Cholinergic modulation

In accordance with [73, 38, 52], we use the fact that, during exploration, high levels of cholinergic modulation are present and the excitatory feedback is thus in part suppressed; during sleep and quiet wakefulness, the ACh levels drop and this suppression is reduced, thus leading to generally stronger synaptic weights. We use a simple 2.5 factor to increase the strength of excitatory synapses during the replay mode in comparison with the encoding mode. Though not too much data is available on the cholinergic effect in vivo, this factor is in line with the one obtained from hippocampal slices of areas CA3 and CA1 in rats [87, 53].

#### 4.2.5 Hebbian plasticity from sequence to place cells

To connect a bistable unit in the sequence encoding module with the corresponding population of place cells, we use a Hebbian like plasticity mechanism. Initially, all synapses from sequence to place cells are silent, and they are potentiated and turned active if the corresponding pre and post cells fire at high rates in a short period. The exact implementation was done by a voltage-based STDP plasticity rule [84] which tends to increase synaptic weight if both pre and post neurons have a high firing rate. As the process of activating a silent synapse is equivalent to LTP [88, 89], we speed up the dynamics to make it consistent with the timescale of LTP. In accordance with [84, 39], the dynamics of the synaptic strength *w_ij_* from sequence cell *j* to place cell *i* are given as:

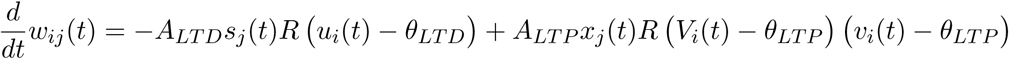

where *R*(*x*) is a linear-rectifying function, that is, *R*(*x*) = 0 if *x ≤* 0 and *R*(*x*) = *x* otherwise; *u_i_* and *v_i_* represent the membrane voltage *V_i_* low-pass filtered with time constants *τ_u_* and *τ_ν_* respectively; and *x_j_* represents the spike train *s_j_* low-pass filtered with time constant *τ_x_*. The constants, *A_LTD_*, *A_LTP_*, *θ_LTD_*, *θ_LTP_ τ_u_*, *τ_v_* and *τ_x_* are given in Table 5 and are the same as [39], with the exception of *A_LTP_*, which is increased to speed up the dynamics and bring the timescale of activating a synapse into the hundred milliseconds range. Furthermore, the maximum value of *w_ij_* is 5*pF*.

**Table 2:**
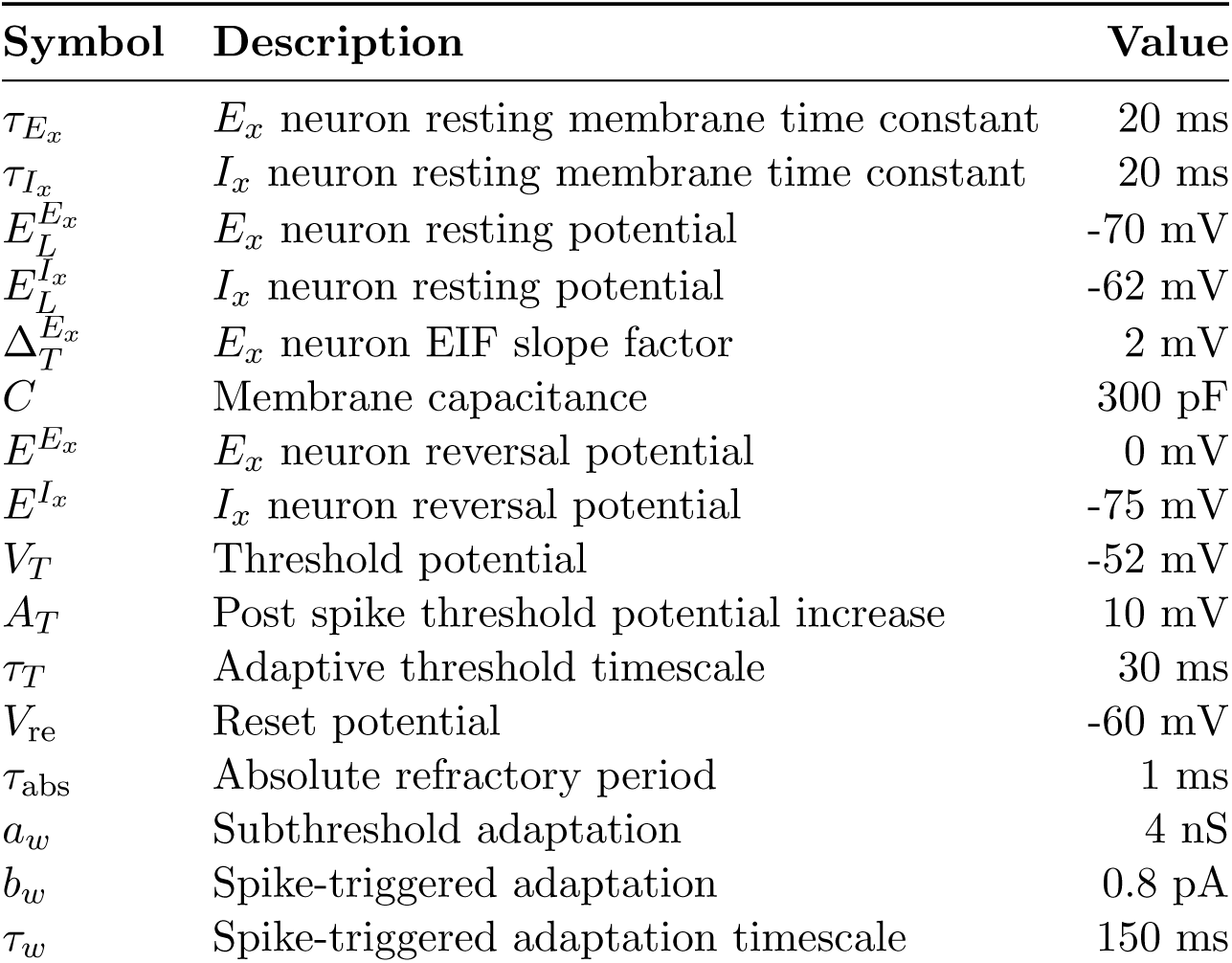
Summary of parameters for membrane dynamics.

### 4.3 Overview of parameters

Here we summarize the parameters of the spiking neural network simulation of exponential integrate-and-fire neurons. Table 2 and Table 3 contain the parameters regarding the membrane potential dynamics of a neuron, and the parameters regarding the neuronal coupling, respectively. Table 4 contains the dendritic spikes parameters that were used, and Table 5 the Hebbian plasticity ones.

For Figure 1, we use a multiplicative gaussian noise 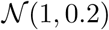 for each individual synaptic weight (shown in Table 3) in the spiking network. For Figure 5C,D, we use a different set of synaptic weights which we show in Table 6.

**Table 3:** Summary of parameters for neuronal coupling.

| Symbol | Description | Value |
| --- | --- | --- |
| $N^{E_x}$ | Number of neurons in population $E_x$ | 200 |
| $N^{I_x}$ | Number of neurons in population $I_x$ | 100 |
| $N^{I_C}$ | Number of neurons in population $I_C$ | 100 |
| $N^{I_G}$ | Number of neurons in population $I_G$ | 100 |
| $p$ | Connection probability | 0.2 |
| $\tau_r^E$ | Rise time for excitatory synapses | 1 ms |
| $\tau_d^E$ | Decay time for excitatory synapses | 6 ms |
| $\tau_r^I$ | Rise time for inhibitory synapses | 0.5 ms |
| $\tau_d^I$ | Decay time for inhibitory synapses | 2 ms |
| $r_{\text{ext}}^{E_x E}$ | Rate of external excitatory input | 1.0 kHz |
| $r_{\text{ext}}^{E_x I}$ | Rate of external inhibitory input | 1.0 kHz |
| $r_{\text{ext}}^{I_x E}$ | Rate of external excitatory input | 1.0 kHz |
| $r_{\text{ext}}^{I_x I}$ | Rate of external inhibitory input | 1.0 kHz |
| $r^{X I_G}$ | Rate of $I_G$ input to $X$ | 0.05 kHz |
| $w^{X I_G}$ | Weight of $I_G$ input to $X$ | 2.5 pF |
| $w^{E_x E_x}$ | Excitatory to excitatory strength | 10 pF |
| $w^{E_x I_x}$ | Inhibitory to excitatory strength | 10 pF |
| $w^{I_x E_x}$ | Excitatory to inhibitory strength | 4 pF |
| $w^{I_x I_x}$ | Excitatory to inhibitory strength | 3 pF |
| $w^{E_{x+1} E_x}$ | Forward afferent strength | 1.6 pF |
| $w^{E_x E_{x+1}}$ | Backward afferent strength | 0.8 pF |
| $w^{I_C E_x}$ | Excitatory to competitive inhibitory strength | 1.5 pF |
| $w^{E_x I_C}$ | Competitive inhibitory to excitatory strength | 12 pF |
| $w_{\text{ext}}^{E_x E}$ | External excitation to excitation strength | 2 pF |
| $w_{\text{ext}}^{I_x E}$ | External excitation to inhibition strength | 0 pF |
| $w_{\text{ext}}^{E_x I}$ | External inhibition to excitation strength | 0 pF |
| $w_{\text{ext}}^{I_x I}$ | External inhibition to inhibition strength | 4 pF |

**Table 4:**
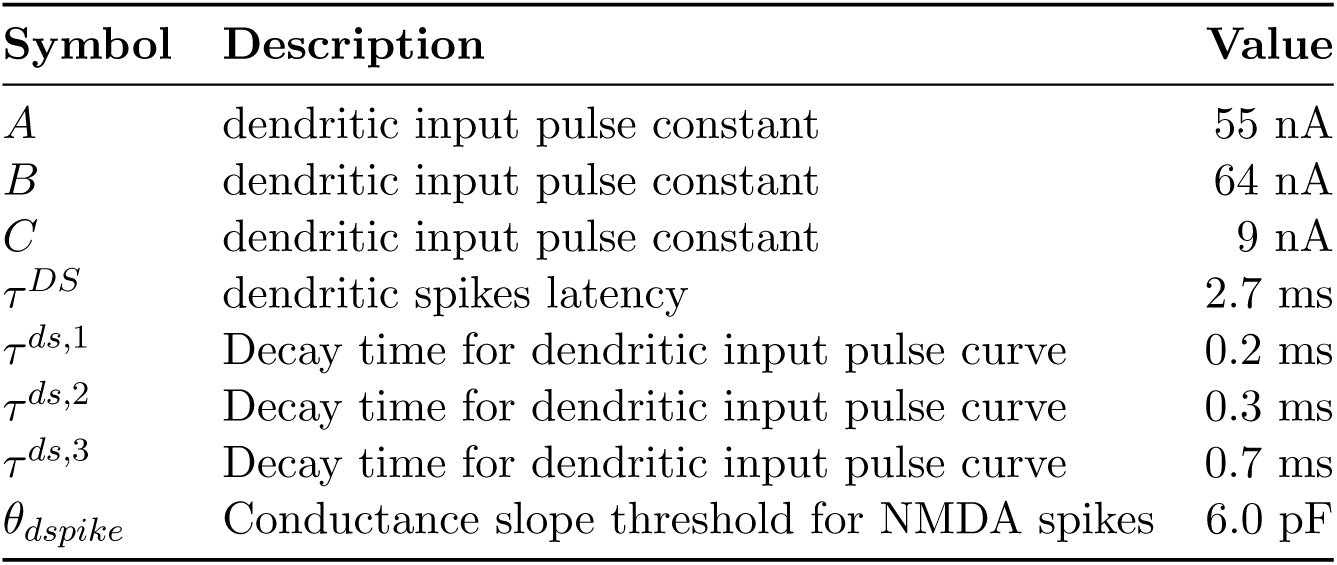
Summary of parameters for dendritic spikes.

**Table 5:** Summary of plasticity parameters.

| Symbol | Description | Value |
| --- | --- | --- |
| $A_{LTD}$ | LTD strength | $0.0008 pAmV^{-1}$ |
| $A_{LTP}$ | LTP strength | $0.07 pAmV^{-1}$ |
| $\theta_{LTD}$ | threshold to recruit LTD | -70 mV |
| $\theta_{LTP}$ | threshold to recruit LTP | -49 mV |
| $\tau_u$ | time constant of low-pass filtered voltage (LTD) | 10 ms |
| $\tau_v$ | time constant of low-pass filtered voltage (LTP) | 7 ms |
| $\tau_x$ | time constant of low-pass filtered spike-train | 15 ms |

**Table 6:** Summary of alternative parameters for neuronal coupling.

| Symbol | Description | Value |
| --- | --- | --- |
| $r^{XI_G}$ | Rate of $I_G$ input to $X$ | 0.05 kHz |
| $w^{XI_G}$ | Weight of $I_G$ input to $X$ | 1.5 pF |
| $w^{E_x E_x}$ | Excitatory to excitatory strength | 8 pF |
| $w^{E_x I_x}$ | Inhibitory to excitatory strength | 6 pF |
| $w^{I_x E_x}$ | Excitatory to inhibitory strength | 4 pF |
| $w^{I_x I_x}$ | Excitatory to inhibitory strength | 2 pF |
| $w^{E_{x+1} E_x}$ | Forward afferent strength | 1.3 pF |
| $w^{E_x E_{x+1}}$ | Backward afferent strength | 0.65 pF |
| $w^{I_C E_x}$ | Excitatory to competitive inhibitory strength | 1.8 pF |
| $w^{E_x I_C}$ | Competitive inhibitory to excitatory strength | 5.5 pF |
| $w_{\text{ext}}^{E_x E}$ | External excitation to excitation strength | 3 pF |
| $w_{\text{ext}}^{I_x E}$ | External excitation to inhibition strength | 0 pF |
| $w_{\text{ext}}^{E_x I}$ | External inhibition to excitation strength | 0 pF |
| $w_{\text{ext}}^{I_x I}$ | External inhibition to inhibition strength | 3 pF |

## 5 Additional information

### 5.2 Author contributions

The project idea came from AS and the analysis was performed in collaboration by all the authors. MMG did all simulations, prepared the figures and the video and wrote a first draft of the manuscript. All authors helped revise the manuscript and everyone approved the final version of it.

### 5.4 Supplementary material

A video illustrating the main ideas and contributions of this paper can be found here: http://www.as.inf.ethz.ch/people/members/mmarcelo/sequencereplay/video.html.

## Footnotes

1 In the model, we do not speculate how place cells could correspond to a particular place in an environment. Instead, when the mouse is at a certain location, we enforce that the corresponding place cell ensemble fires at the correct rate, as is experimentally observed [24].

2 In the model, we follow the same approach as [47, 48, 37] and assume that given enough synchronous input dendritic spikes will occur.

3 Lowering the stable rate at which a bistable unit is active would also have undesirable consequences as one needs to be able to clearly distinguish between the low and high firing rate stable points of the bistable units. This will be further discussed in the next subsection.

